## Supplementary figures for "Control of tongue movements by the Purkinje cells of the cerebellum"

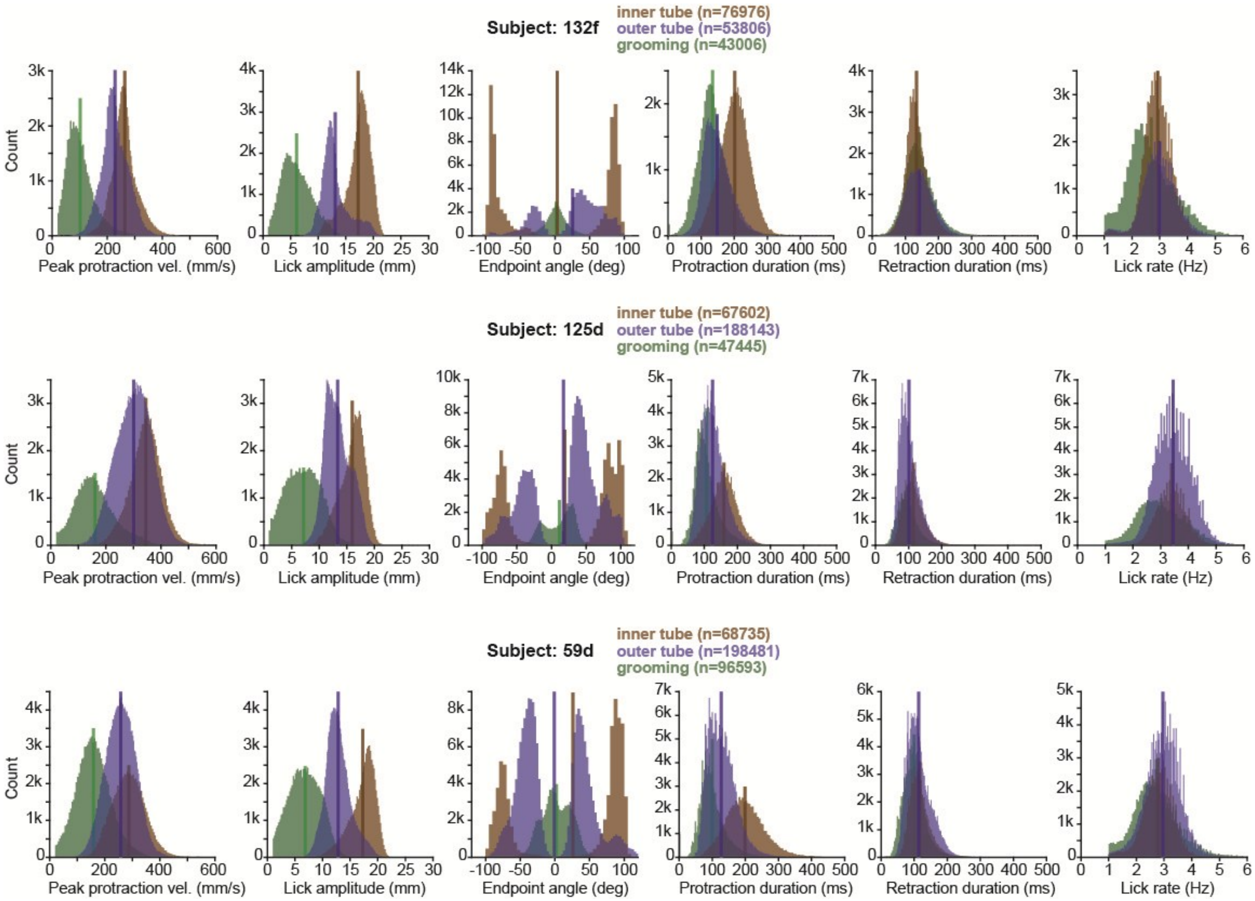


**Supplementary Fig. S1. The kinematic properties of licks in each monkey.**

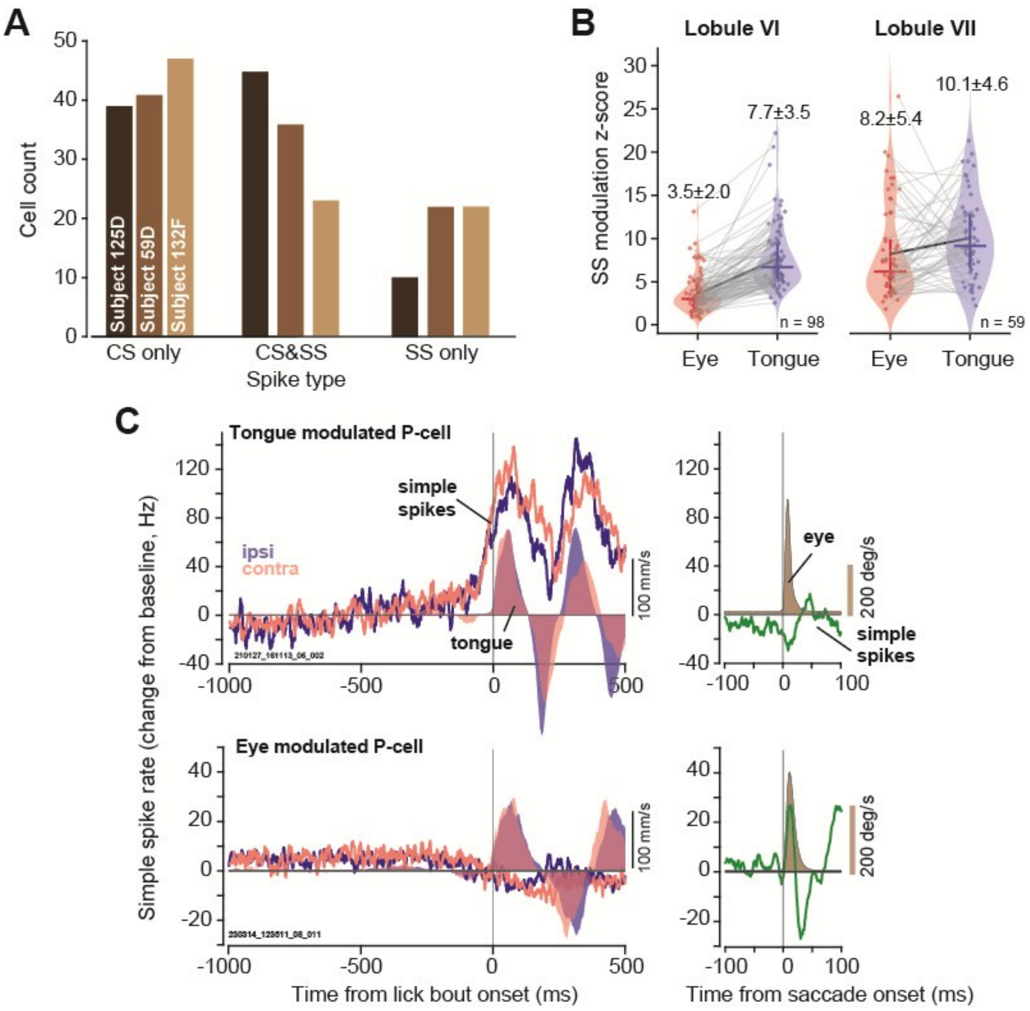


**Supplementary Fig. S2**. **Properties of the P-cells in the database. A.** The number of neurons recorded from each marmoset. CS only refers to P-cells for which only the complex spikes were recorded. CS&SS refers to P-cells for which both the CS and SS were recorded. SS only refers to putative P-cells for which only the SS were recorded. **B.** Modulation index of each P-cell during licking and saccades. The values indicate the mean±SD of each distribution. **C.** Simple spike modulation in two example P-cells. The activities in each cell are aligned to lick bout onset, and saccade onset, both for reward relevant movements.

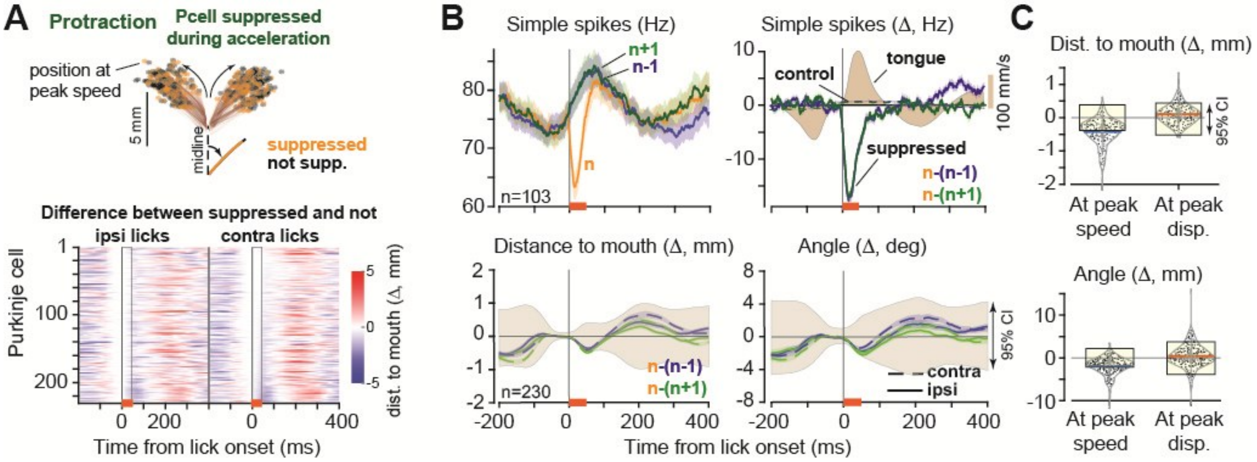


**Supplementary Fig. S3. CS-induced SS suppression did not affect movements during protraction acceleration period. A**. Suppression took place during the acceleration period of protraction. Traces show average tongue trajectory during the acceleration period of protraction for each P-cell during suppressed and control licks. Ipsilateral licks are shown to the left and contralateral to the right. Heatmap quantifies change in endpoint trajectory between suppressed and control licks for each cell. Period of suppression is indicated by the orange bar at the bottom of the heatmap. **B**. Top row: SS rates for licks $\{n-1, n, n+1\}$, where only lick $n$ experienced a CS. Filled color curves indicate tongue velocity. Second row: trajectory of the tongue in lick $n$ as compared to its two temporally neighboring licks. Trajectory is measured via distance from tip of the tongue to the mouth and angle of the tip with respect to midline. The filled region is 95% CI. **C**. Distance to mouth and angle in lick $n$ as compared to neighboring licks. Shaded region is 95% CI.

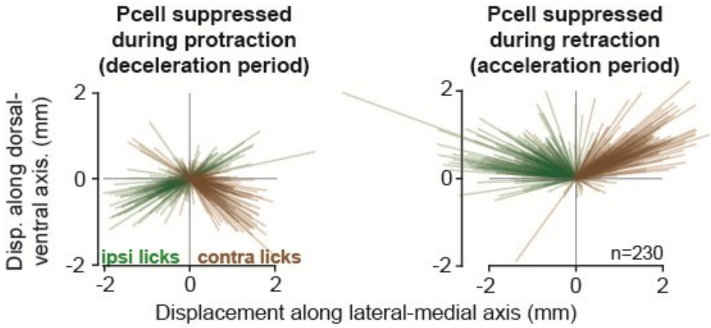


**Supplementary Fig. S4**. **Kinematic effects of CS-induced SS suppression in Cartesian coordinates.** Left plot shows the change in the position of the tip of the tongue at the end of protraction in movements that experienced a CS during the deceleration period of protraction, with respect to movements that did not experience a CS during any period. Right plot shows the change in position of the tip of the tongue as measured at peak retraction velocity, comparing movements that experienced a CS during the retraction acceleration period with movements that did not experience a CS in any period. Each line is the average effect for a single P-cell. P-cell suppression during protraction produced hypermetria, whereas the suppression during retraction resulted in slowing.

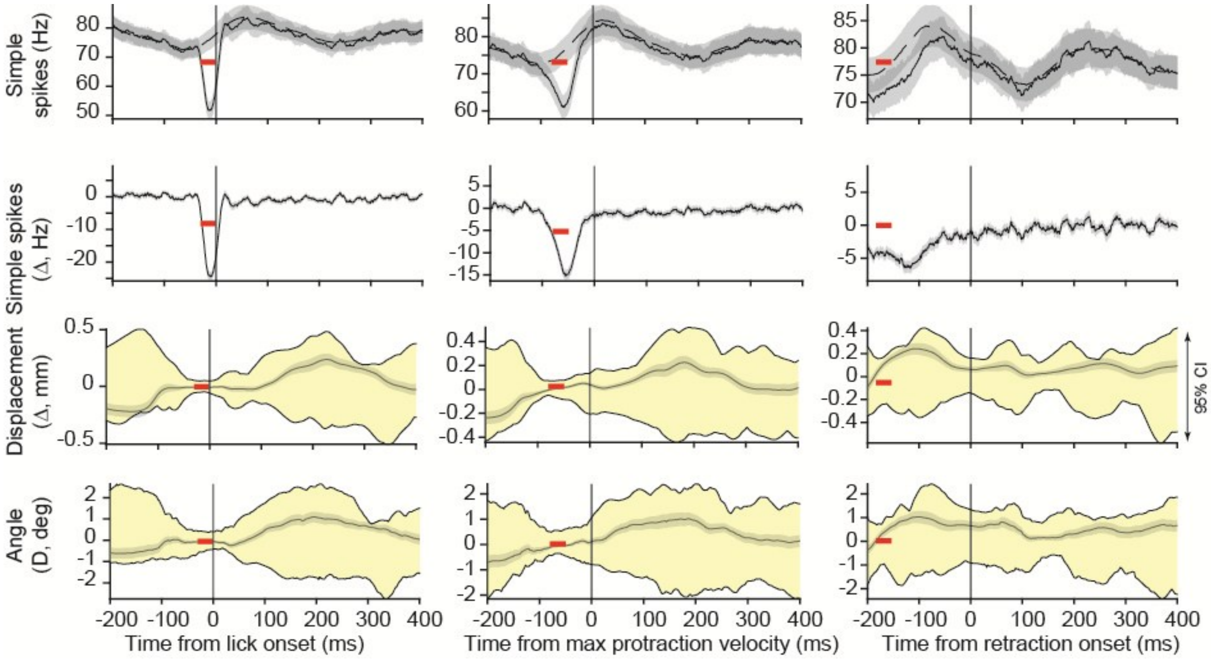


**Supplementary Fig. S5**. **There was little or no change in lick kinematics when the CS-induced SS suppression occurred before lick onset.** We chose the period following completion of one lick and the start of the next lick, which was on average 30ms in duration, i.e., the period in which the tongue was in the mouth. Top row: SS rates, aligned to lick onset, protraction peak speed, and retraction onset. Second row: change in SS rates. Third row: change in tongue displacement. Fourth row: change in tongue angle. The filled region is 95% confidence interval. Error bars are SEM.

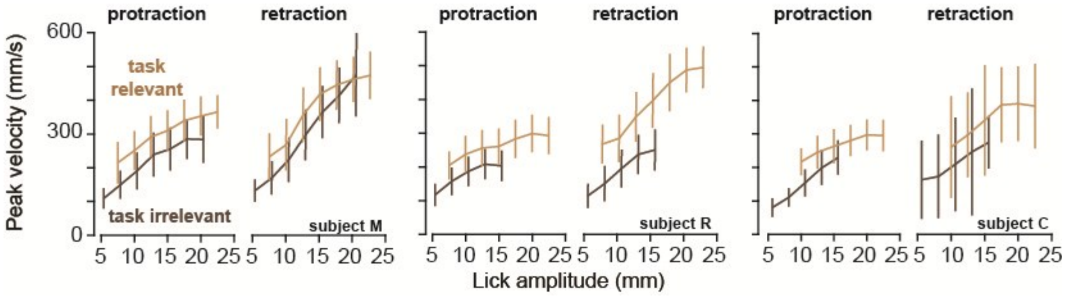


**Supplementary Fig. S6**. **Peak velocity as a function of lick amplitude during protraction and retraction for task relevant and task irrelevant licks in each subject.** Error bars are SEM.

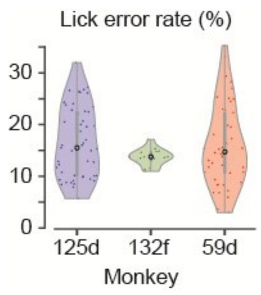


**Supplementary Fig. S7**. **Percent of licks in which the food was inside the tube, but the tongue missed the tube's entrance and did not contact the food**.

**Supplementary Table 1. Distribution of tongue and eye modulated P-cells in the vermis region of the cerebellum**

| Region | Eye Modulated | Tongue Modulated | Only Eye Modulated | Only Tongue Modulated | Both | Neither | Saccade z-score (mean ± SD) | Lick z-score (mean ± SD) |
|---|---|---|---|---|---|---|---|---|
| **Lobule VI** | 49/98 (50.00%) | 97/98 (98.98%) | 0/98 (0.00%) | 48/98 (48.98%) | 49/98 (50.00%) | 1/98 (1.02%) | 3.52 ± 2.05 | 7.67 ± 3.52 |
| **Lobule VII** | 56/59 (94.92%) | 57/59 (96.61%) | 2/59 (3.39%) | 3/59 (5.08%) | 54/59 (91.53%) | 0/59 (0.00%) | 8.21 ± 5.39 | 10.06 ± 4.64 |
