## Supplementary figures and images for "Control of tongue movements by the Purkinje cells of the cerebellum"

### video1 (lick bout with error).gif

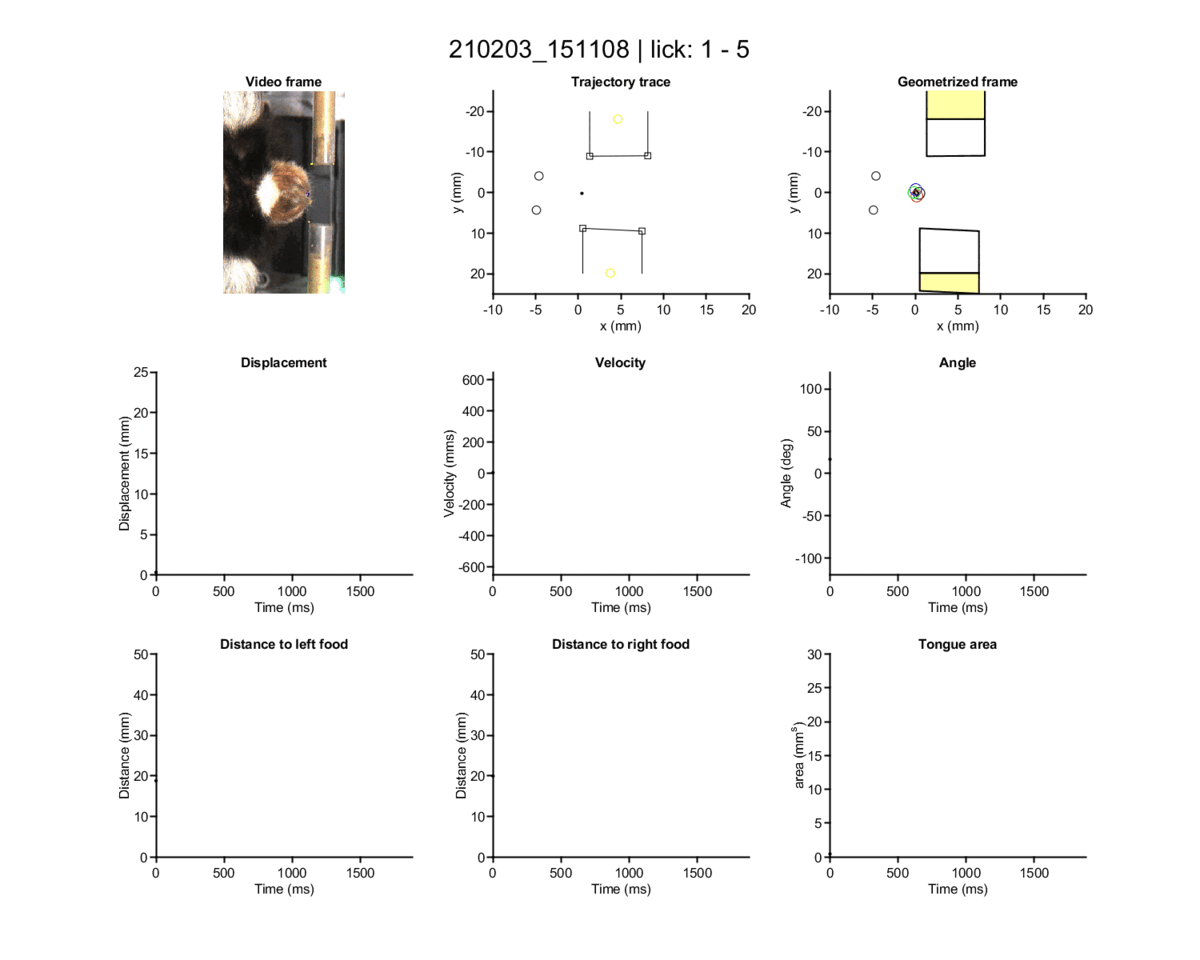

### video 2 (grooming).gif

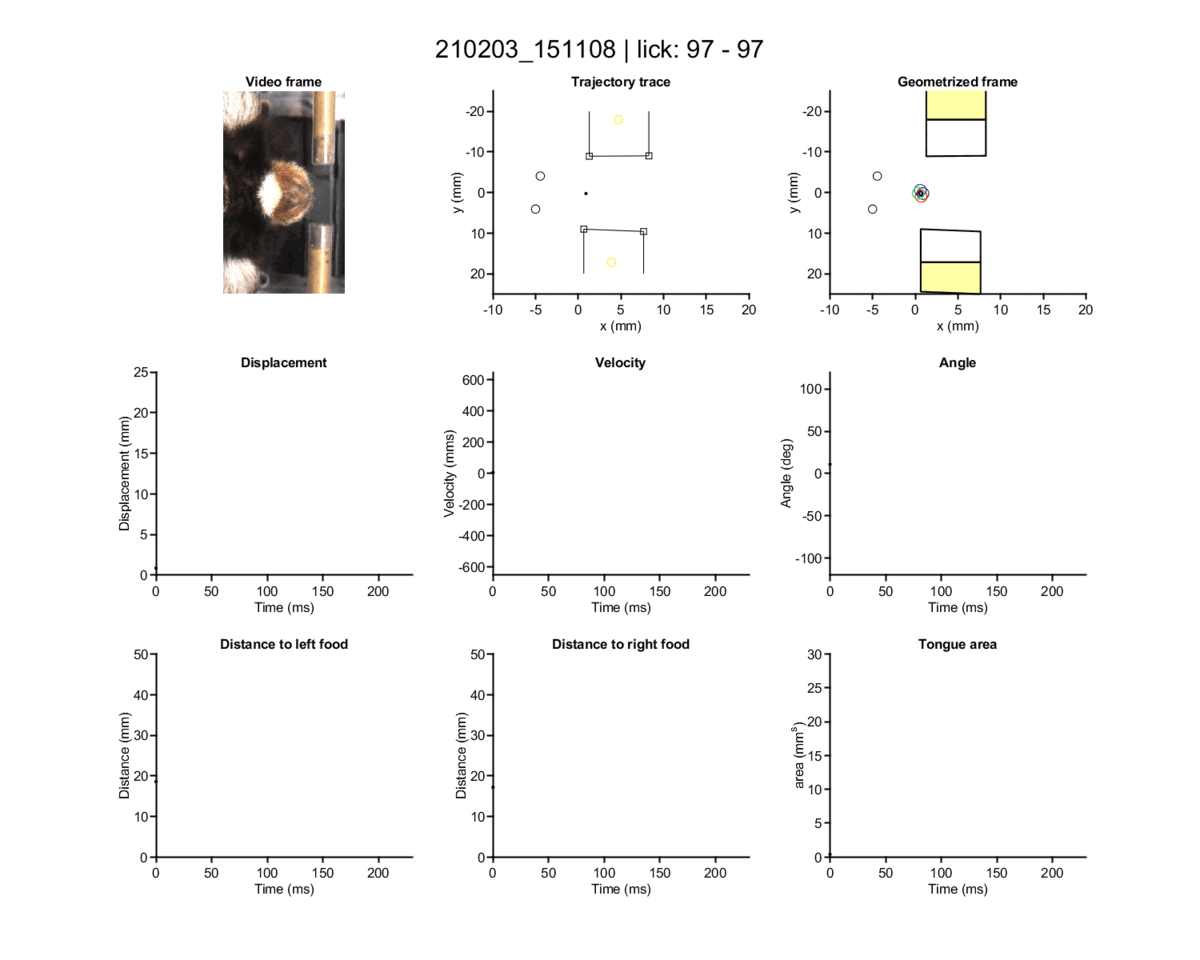

### video 3 (grooming).gif

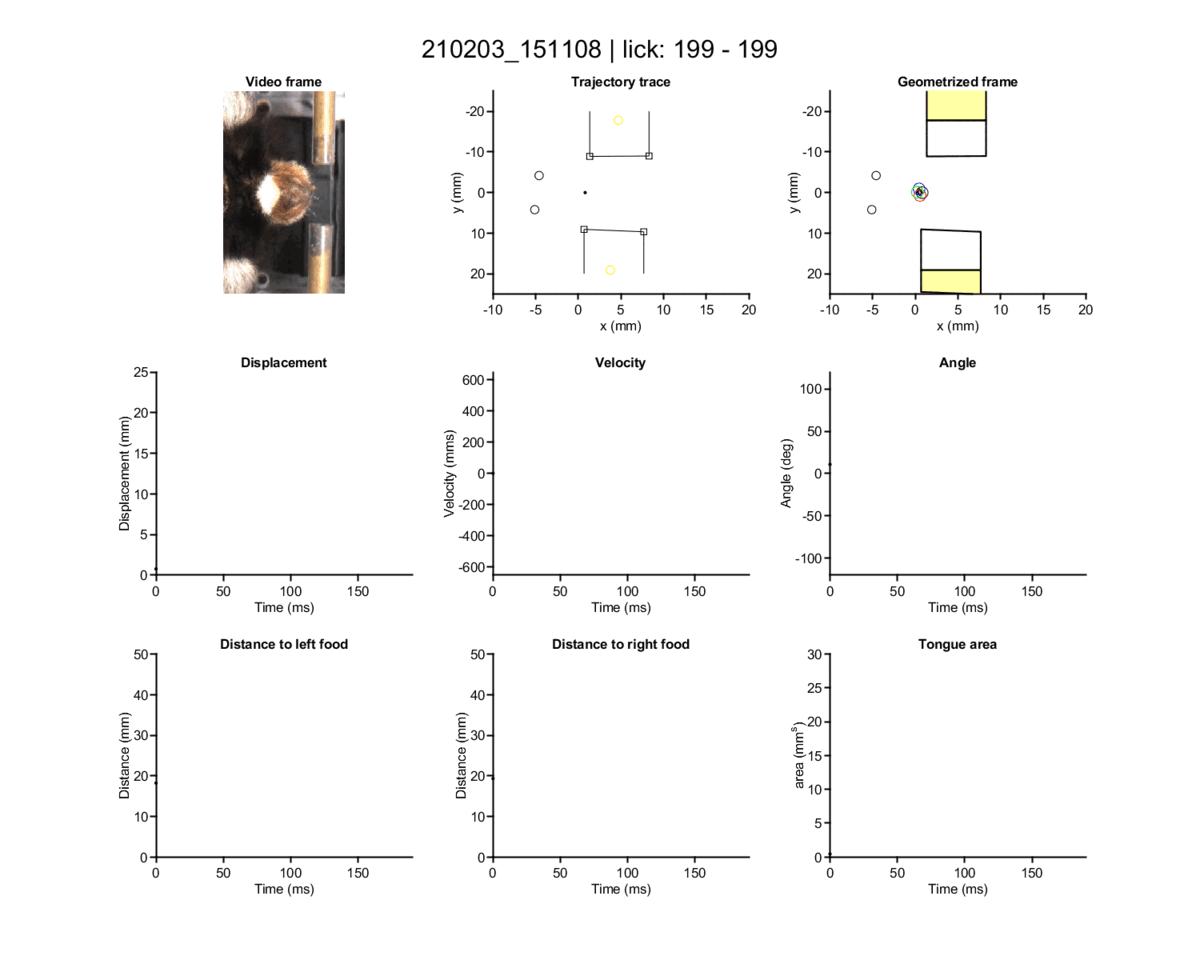

### video 4 (grooming bout).gif

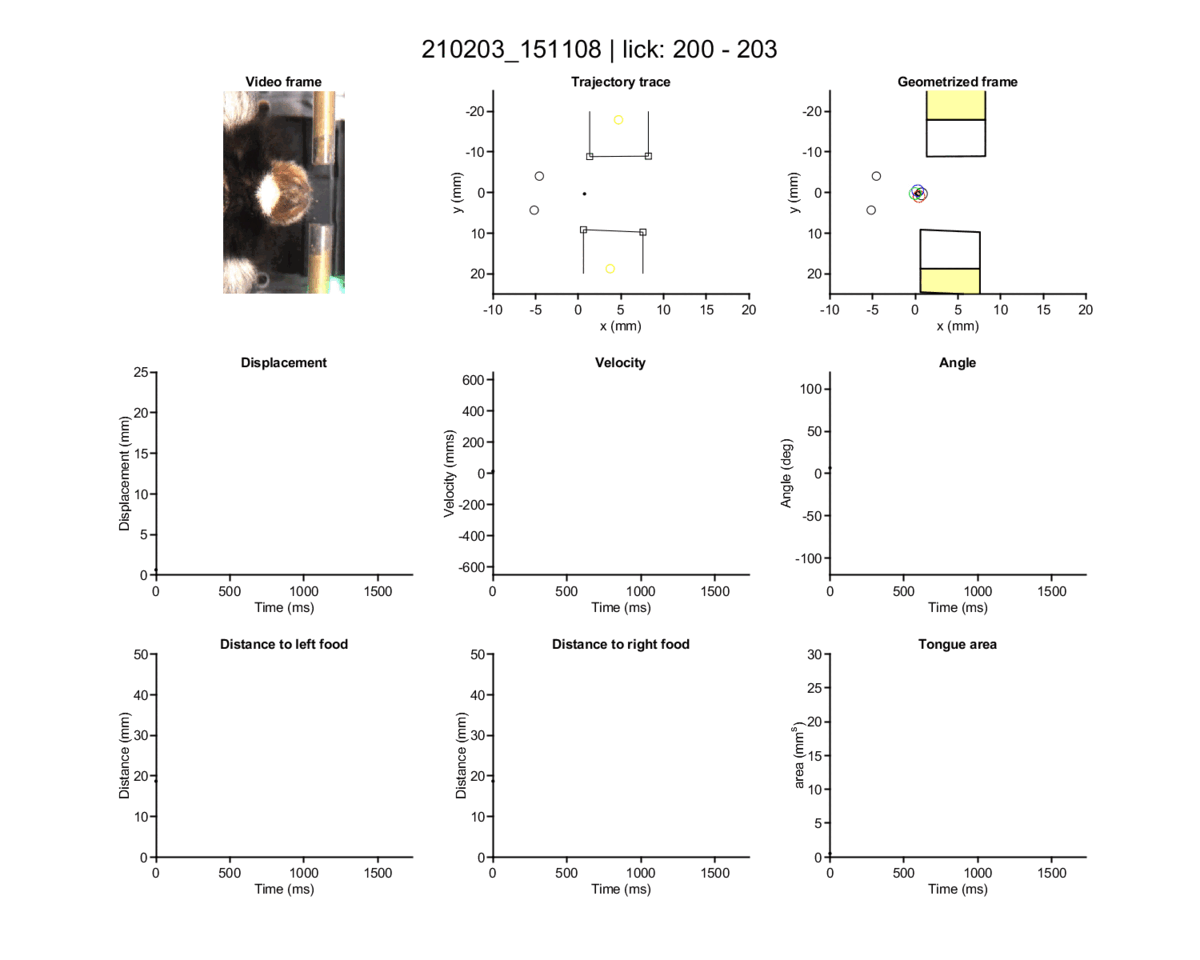

### video 5 (outer tube).gif

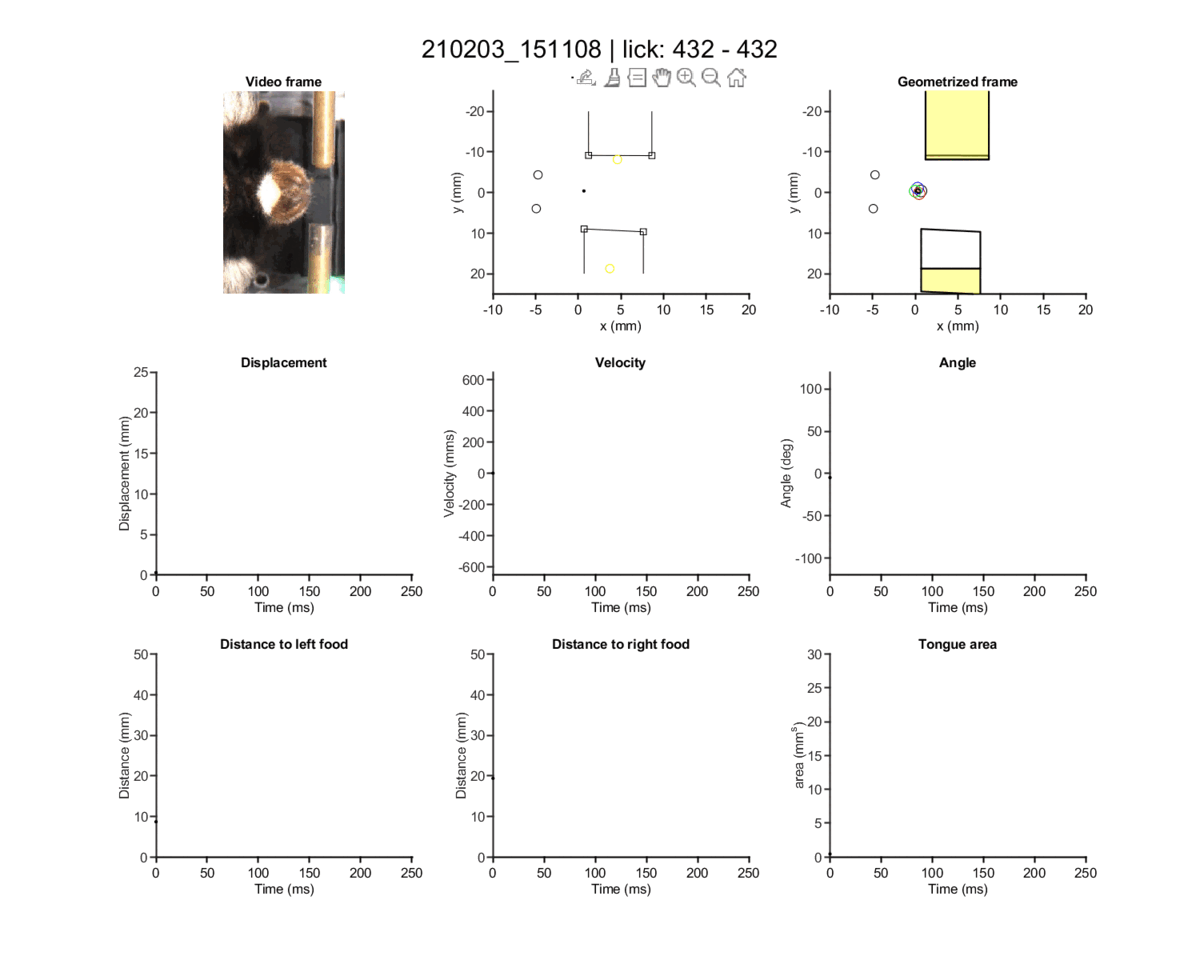

### video 6 (inner tube 1).gif

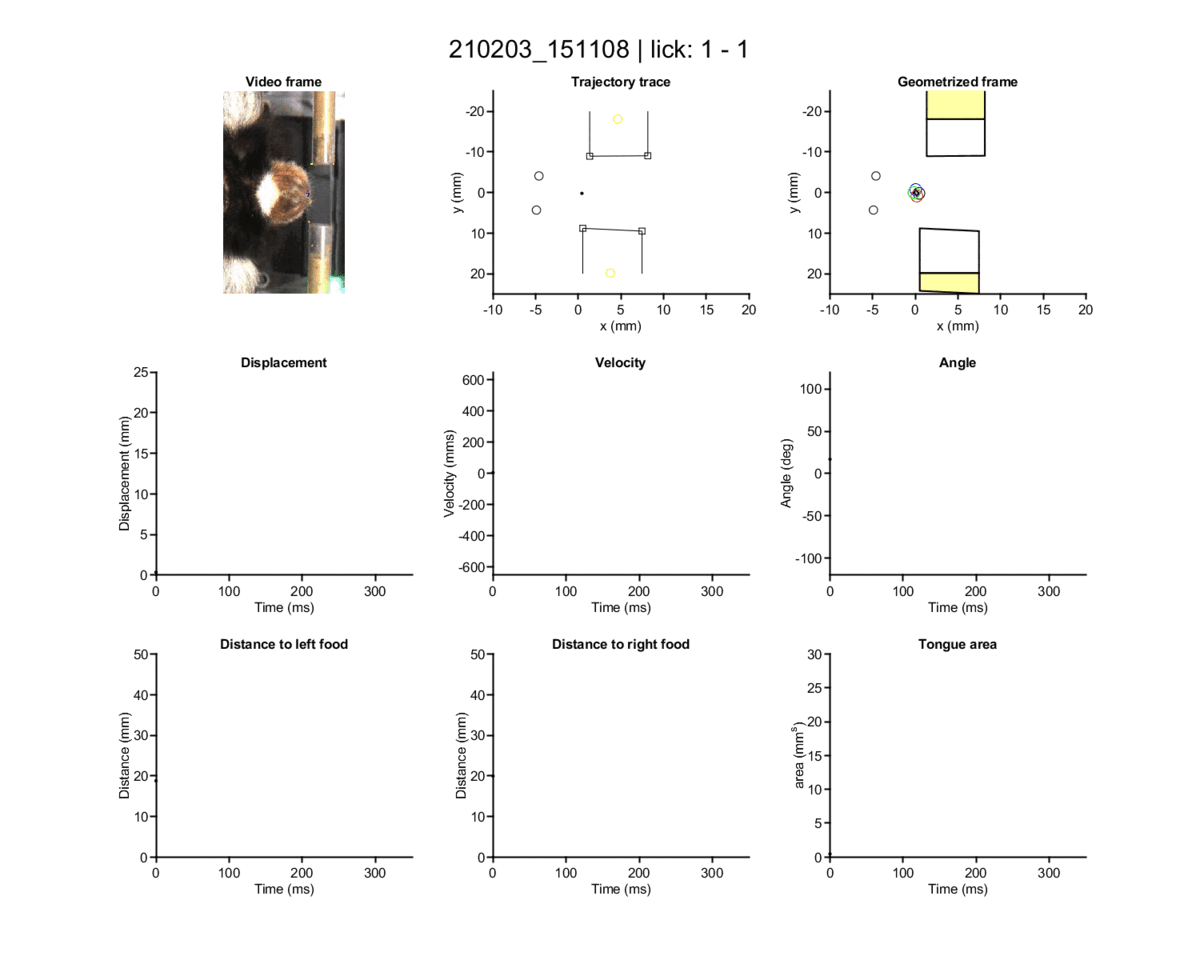

### video 7 (inner tube 2).gif

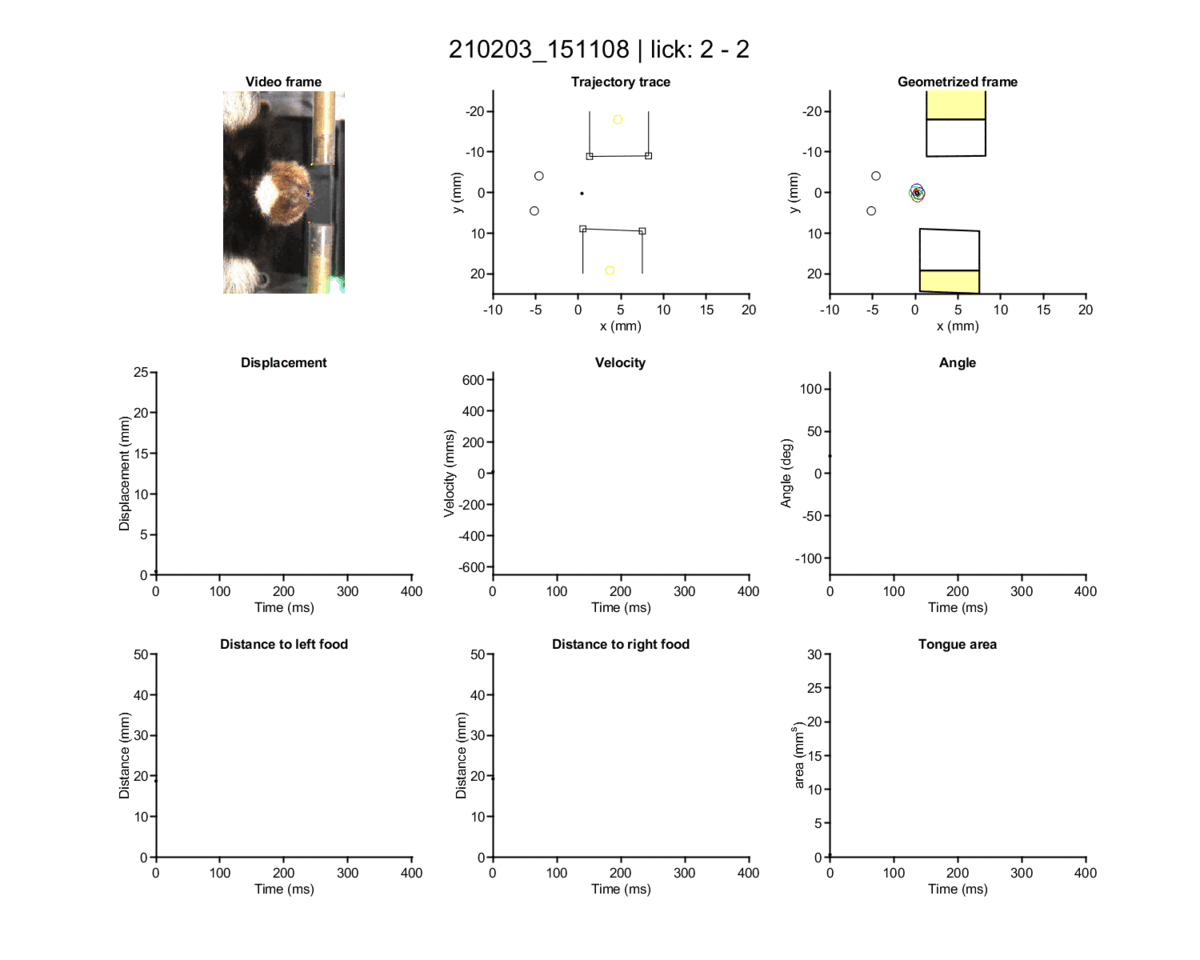

### video 8 (outer tube error).gif

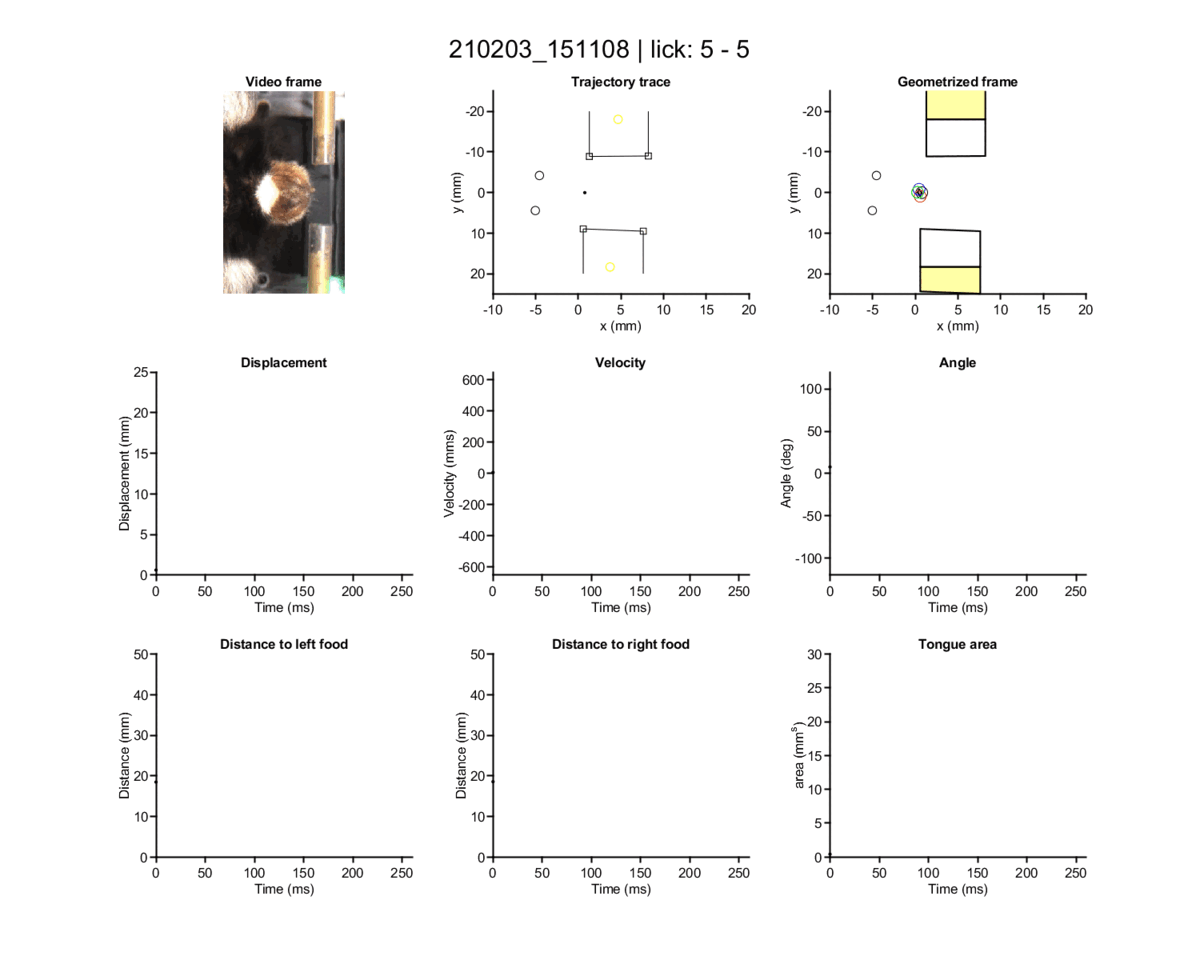

### video 9 (outer tube error).gif

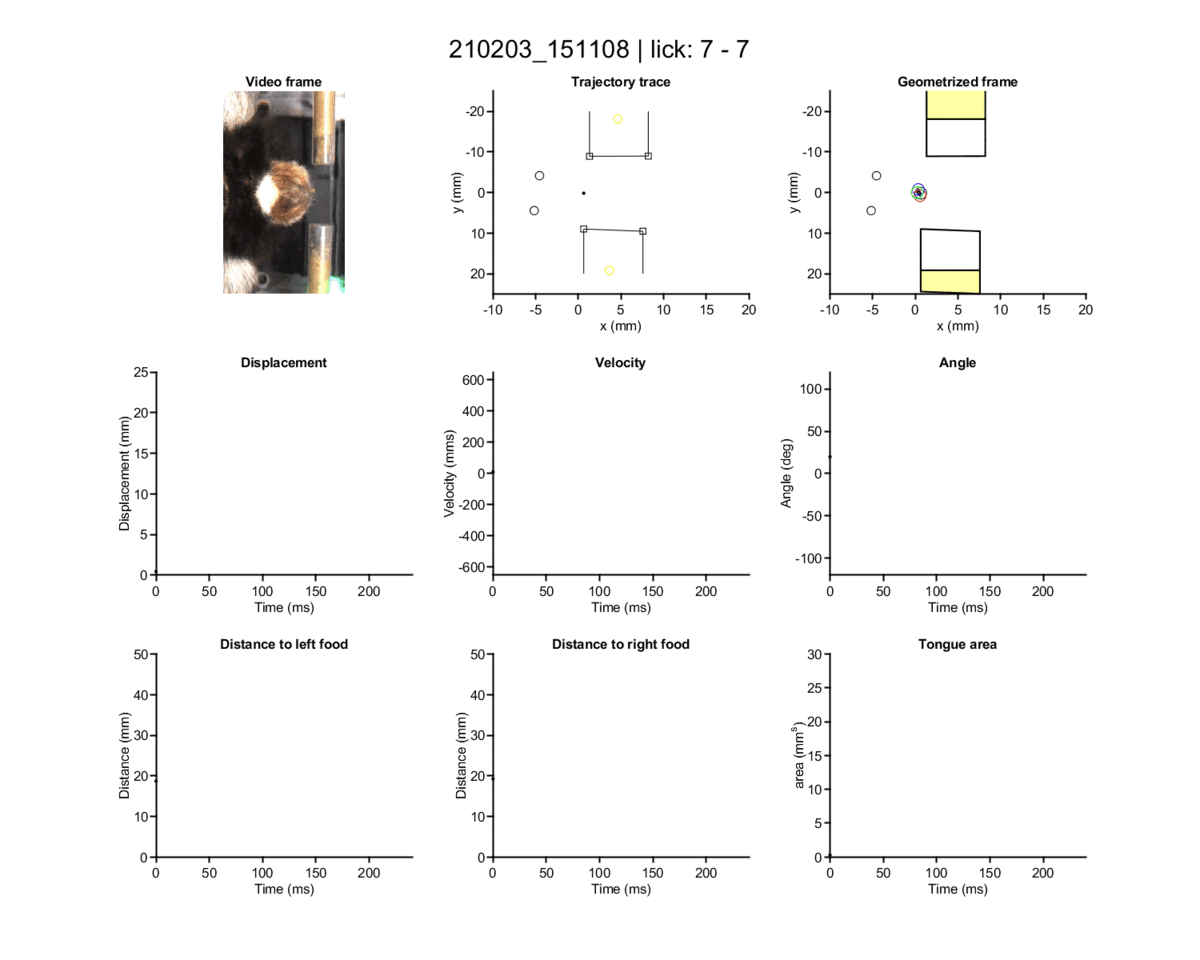

### video 10 (outer tube error).gif

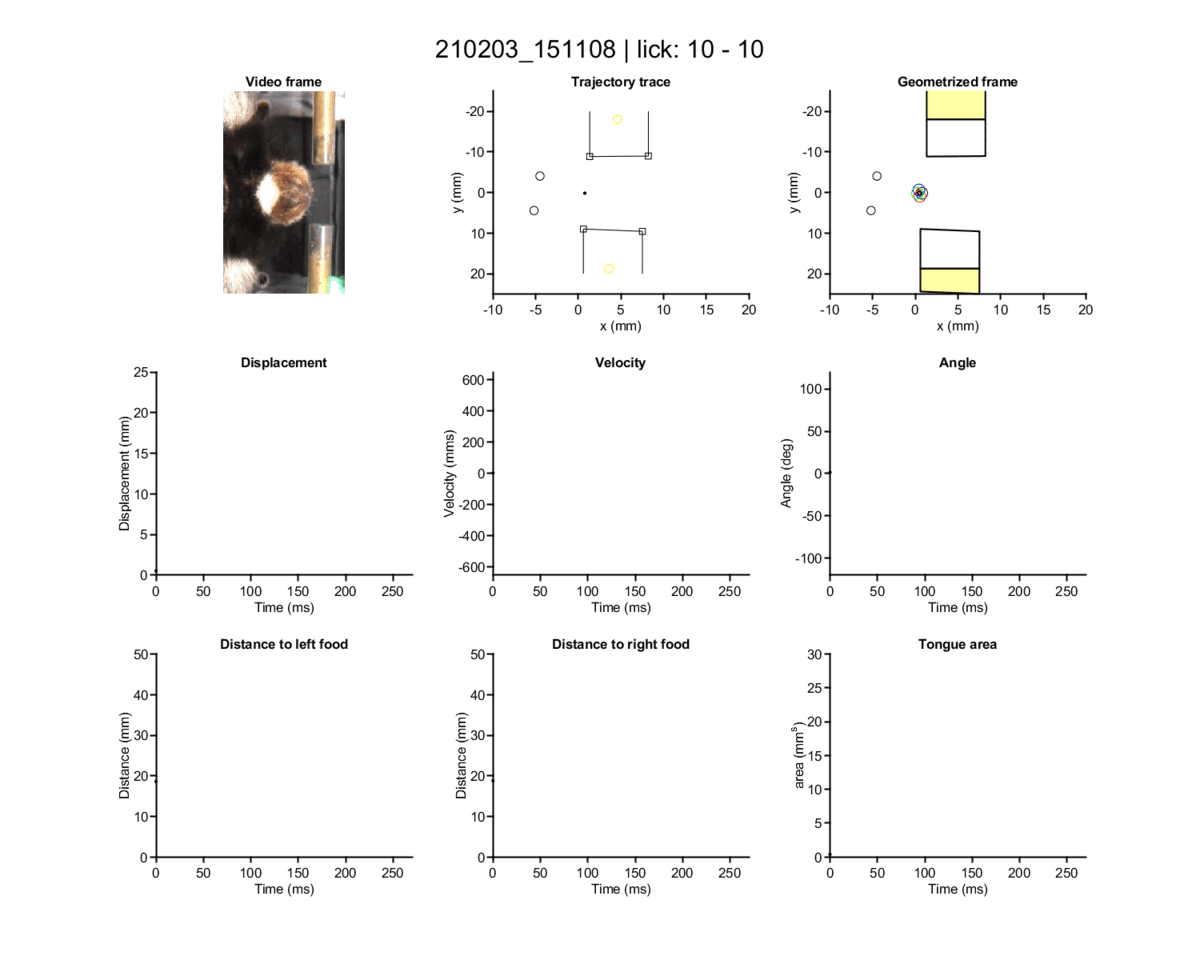
